# Potato Agent: AI-Driven Data and Knowledge Exploration on an Agent-Ready Potato Multi-Omics Platform

**DOI:** 10.64898/2026.08.12.744101

**Authors:** Yarui Dong, Jinye Li, Futing Li, Jilin Luo, Yudong Jia, Dawei Li, Li Wang, Xiaoqi Su, Juan Hu, Yi Shang, Sanwen Huang, Yujuan Zhu, Yuxin Jia

## Abstract

Potato is an important non-cereal food crop worldwide. However, the limited number of functionally validated genes remains a major bottleneck to favorable allele stacking and genome design breeding in potato. Rapid advances in AI agents offer a promising means to support crop breeding by translating natural-language questions into coordinated data analysis and knowledge retrieval. Their reliable use for potato breeding, however, is constrained by fragmented multi-omics resources that lack consistent curation and machine-accessible interfaces. Here, we constructed an agent-ready potato multi-omics database integrating genomic resources from 150 potato accessions, 259 bulk RNA-seq samples, and 14 spatial transcriptomic datasets into a pangenome, a tissue expression atlas, co-expression networks, and spatial expression maps accessible through open APIs. We developed 39 potato-specific Agent Skills for reproducible bioinformatics analysis and comprehensive data and knowledge exploration, enabling natural-language questions to be translated into standardized data-retrieval and analysis tasks. By integrating direct evidence from potato studies, functions of homologous genes in Arabidopsis, rice, and maize, and tissue expression patterns, we generated genome-wide functional predictions for 37,658 genes in the DM reference genome. We further developed Potato Agent as a multi-user, browser-based platform with isolated workspaces and online result preview, reducing the technical burden of agent deployment and providing direct access to integrated data, knowledge, and workflows. Case studies demonstrated its capabilities in reproducible bioinformatics analysis, agent-assisted identification of a tuber development regulator, scientific data visualization, and haplotype-aware promoter analysis and sgRNA design. Together, the agent-ready database and Potato Agent provide an integrated infrastructure for functional gene discovery and hybrid breeding in potato.

## Introduction

Potato is an important non-cereal food crop worldwide. In recent years, diploid hybrid breeding has offered a new route to overcome the constraints imposed by polyploidy and high heterozygosity in conventional tetraploid breeding and to accelerate genetic gain (Lindhout et al., 2011; Zhang et al., 2021). However, both diploid hybrid breeding and conventional tetraploid breeding depend on a detailed understanding of the genetic basis of key agronomic traits. The number of potato genes that have been cloned and functionally validated remains limited, constraining the efficient identification and precise stacking of favorable alleles and the implementation of molecular design breeding. This knowledge gap has become a shared bottleneck for both breeding systems (Gebhardt, 2023).

Well-developed biological databases are important infrastructure for functional gene discovery. The core content of resources such as TAIR, MaizeGDB, RiceData, and RAP-DB can be broadly grouped into two categories. The first comprises gene structures, functional annotations, mutants or phenotypes, and associated publications, allowing users to retrieve existing evidence starting from a gene or trait. The second comprises multi-omics data, including genomes, genetic variation, and transcriptomes, which can be used for gene mapping, expression comparisons, and candidate gene screening. Integrating these two types of resources connects genes with experimental evidence and agronomic traits and can facilitate gene function studies and molecular breeding (Andorf et al., 2016; Reiser et al., 2024; Sakai et al., 2013).

To address the fragmented nature of potato gene function information, inconsistent gene identifiers, and inefficient literature retrieval, we developed and released the Potato Knowledge Hub in 2025 (Li et al., 2026). This resource systematically organizes potato publications and functional genes and supports natural-language question answering, gene searches, and sequence extraction. Existing potato data platforms, including the Potato Knowledge Hub and Spud DB, have generally emphasized either literature and functional gene curation or selected genomic resources (Hirsch et al., 2014). The unified integration of functional genes, supporting publications, and multi-omics data remains limited. In addition, many conventional multi-omics databases rely on predefined datasets, query interfaces, and analysis modules. This structure can make it difficult for users to select data, combine analysis steps, and construct reproducible workflows tailored to a specific scientific question, thereby limiting more flexible use of the available resources.

AI agent technology has advanced rapidly in recent years. By querying databases and invoking bioinformatics tools, an agent can translate a natural-language question into a sequence of data queries, analyses, and evidence integration, potentially supporting greater automation in crop gene discovery (Jin et al., 2024; Zhou et al., 2024). Reliable execution, however, depends on data resources designed for machine access. Here, we define agent-ready data as resources with unified identifiers, explicit reference versions, DOI-based provenance records, machine-readable data structures, and stable application programming interfaces (APIs). Potato multi-omics data are currently distributed across different sources and often lack consistent curation, cleaning, and calibration of file formats, metadata, and reference genome versions. These limitations hinder data integration and reproducible analysis. Agent deployment also requires environment configuration and tool integration, creating a technical barrier for breeders and wet-laboratory researchers.

In this study, we constructed an agent-ready potato multi-omics database that integrates 150 potato accessions, including 149 publicly available genome resources and a newly assembled, highly heterozygous diploid accession, 01-58. We also collected 259 bulk RNA-seq samples and 14 spatial transcriptomic datasets to establish structured resources, including a pangenome, a gene expression atlas, and co-expression networks, all accessible to agents through open APIs. To support data and knowledge use, we packaged database queries, literature retrieval, sequence extraction, and common bioinformatics workflows as specialized Agent Skills. Validated Snakemake workflows were used to improve the consistency and reproducibility of analyses. We further integrated direct evidence from potato studies, functions of homologous genes in Arabidopsis, rice, and maize, and tissue expression patterns to establish a genome-wide functional prediction and evidence-grading system. Finally, we developed Potato Agent, which supports Linux user-based data separation and online result preview, to reduce the technical burden of agent deployment and use. We assessed the practical utility of the system through case studies involving BSA-seq analysis, developmental regulator identification, scientific visualization, as well as promoter analysis and sgRNA design.

## Results

### Construction of an agent-ready potato multi-omics data foundation

The potato genomic and transcriptomic data accumulated in recent years have provided an important basis for investigating species origins, tuber formation, domestication, and genetic variation relevant to hybrid breeding (Cheng et al., 2025; Tang et al., 2022; Zhang et al., 2025). However, these data are distributed across studies and databases and lack consistent standards for file formats, metadata descriptions, and reference genome versions. Consequently, they are difficult for researchers to integrate directly and for AI agents to access reliably. We therefore systematically collected and standardized potato genome, bulk RNA-seq, and spatial transcriptomic data. Each resource was organized with unified identifiers, explicit versions, provenance information, and machine-readable data structures. We then constructed a pangenome, tissue expression atlas, co-expression networks, and spatial expression maps and provided stable, open APIs for unified data access. Together, these resources provide a foundation for agent-driven data retrieval and analysis.

We first collected and curated high-quality public genome resources for 149 potato accessions, comprising 135 diploid accessions and 14 haplotype-resolved tetraploid accessions. Among the diploid accessions, 60 were represented by haploid-level assemblies and 75 by haplotype-resolved assemblies. All 135 diploid accessions were used for pangenome analysis, comprising 90 wild and 45 cultivated potatoes (Figure 1A). Homologous gene clustering identified 2,107 core orthogroups, defined as gene families present in all accessions; 8,510 soft-core orthogroups, present in 122-134 accessions; 189,934 dispensable orthogroups, present in 2-121 accessions; and 3,324 private orthogroups, detected in only one accession (Figure 1A). These results describe extensive variation in gene content among potato germplasm and provide a structured pangenome resource for comparative genomic analyses and candidate gene screening.

**Figure 1.**
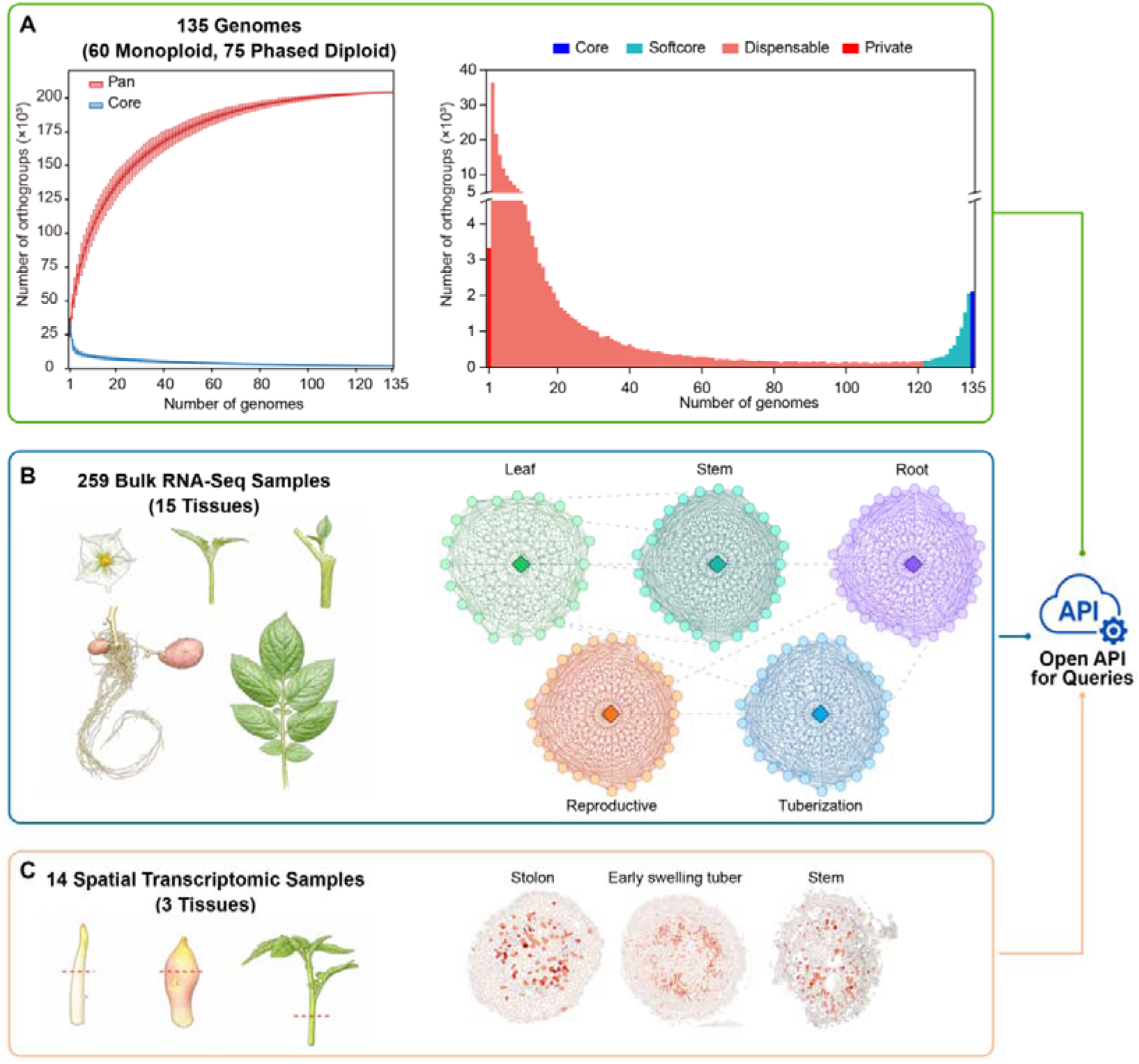
Agent-ready potato multi-omics resources and open API access. (A) Pangenome analysis of 135 diploid potato accessions, including 90 wild and 45 cultivated accessions represented by 60 monoploid and 75 phased diploid assemblies. (B) Tissue expression atlas constructed from 259 bulk RNA-seq samples covering 15 tissue categories, together with co-expression networks for leaf, stem, root, reproductive, and tuberization-related samples. (C) Spatial transcriptomic resources comprising 14 samples from stolons, early swelling tubers, and stems. Red dashed lines indicate the positions of transverse sections. The pangenome, expression, co-expression, and spatial transcriptomic resources are accessible through open APIs.

In addition to the public genomes, we generated a haplotype-resolved genome assembly for the highly heterozygous diploid accession 01-58 to obtain accurate, accession-specific sequences for genetic transformation and genome editing. The estimated genome heterozygosity was 1.93%, indicating a high level of heterozygosity. Using 45.6 Gbp of PacBio HiFi data and 94.5 Gbp of Hi-C data, we obtained two haplotype assemblies of 748.2 Mbp and 753.0 Mbp for haplotypes 1 and 2, respectively (Supplementary Figure 1A, B). Numerous structural variants were detected between the two haplotypes and between each haplotype and the DM reference genome, further illustrating the complex genomic composition of 01-58 (Supplementary Figure 2). By integrating de novo prediction, transcriptomic evidence, and homologous protein evidence, we annotated 34,281 and 32,900 protein-coding genes in the two haplotypes, with BUSCO completeness values of 99.2% and 99.0%, respectively (Supplementary Figure 1C). The genome was incorporated into the platform, enabling researchers and agents to directly retrieve accession-specific gene, promoter, and target-site sequences for downstream functional validation and molecular design.

To establish a tissue expression reference for gene function inference, we collected and uniformly processed 259 bulk RNA-seq samples from public databases (Figure 1B). These samples covered 15 tissue categories, including fruit, anther, perianth, stem, leaf, root, stolon, and tubers at different developmental stages. All datasets were aligned to the DM reference genome using a consistent pipeline, followed by expression quantification and normalization to construct a genome-wide tissue expression atlas. Based on the tissue and developmental attributes of the samples, we further constructed five co-expression networks: leaf, stem, root, reproductive, and tuberization. The standardized expression matrices and co-expression relationships allow the platform to provide evidence for candidate gene screening from both tissue-specific expression and expression association.

We also integrated 14 spatial transcriptomic datasets from stems, stolons, and early swelling tubers generated in our previous work (Figure 1C). Because the raw data were not readily suited to direct querying or cross-sample comparison, we standardized the spatial coordinates, tissue-region annotations, and gene expression matrices and developed an interactive visualization and gene-query interface. Users can examine the spatial expression patterns of target genes in tissues associated with tuber formation, while agents can retrieve the corresponding tissue regions and expression information through standardized interfaces. The resulting spatial expression maps complement the lack of spatial resolution in bulk RNA-seq and provide a basis for investigating cell type- and region-associated regulation during tuber initiation and swelling.

In summary, through unified curation, standardized analysis, and structured storage, we established a potato multi-omics data foundation encompassing genome assemblies, a pangenome, bulk RNA-seq, and spatial transcriptomics, together with open and unified query interfaces. These agent-ready resources support structured retrieval, association analyses, and evidence integration across multiple omics layers and provide a data foundation for potato data exploration, functional gene discovery, and molecular breeding research in an agent-enabled setting.

### Development and optimization of Agent Skills for potato data and knowledge exploration

To enable AI agents to access potato multi-omics data reliably and perform specialized analyses, we developed Agent Skills for potato data and knowledge exploration. Agent Skills package the domain knowledge, procedures, analytical tools, and reference resources required for a specific task into functional units that can be called as needed, thereby improving the consistency, reusability, and reliability of agent execution. Although open-source projects such as scientific-agent-skills (https://github.com/K-Dense-AI/scientific-agent-skills) and bioSkills (https://github.com/GPTomics/bioSkills) provide skills for a range of bioinformatics tasks, most existing skills address general analytical settings and have not been specifically adapted to the high heterozygosity of potato genomes or the diversity of potato reference genomes and gene identifier systems.

We therefore developed and optimized 39 Agent Skills for potato research on the basis of analytical methods validated in published studies. These comprised 24 bioinformatics analysis skills and 15 data and knowledge query skills (Figure 2). All skills are openly available, and their task instructions, analysis scripts, workflows, and configuration files can be directly accessed, reused, and extended by researchers (https://github.com/biojiayuxin/potato-agent/tree/lite/skills/potato-knowledge-bioinfor matics). The bioinformatics skills cover sequence processing and common multi-omics analyses, whereas the data and knowledge query skills support retrieval of publications, functional genes, homologous genes, sequences, and multi-omics data. With this design, an agent can select appropriate skills for a scientific question and translate natural-language instructions into standardized data-retrieval or analysis tasks without generating an entire workflow from scratch.

**Figure 2.**
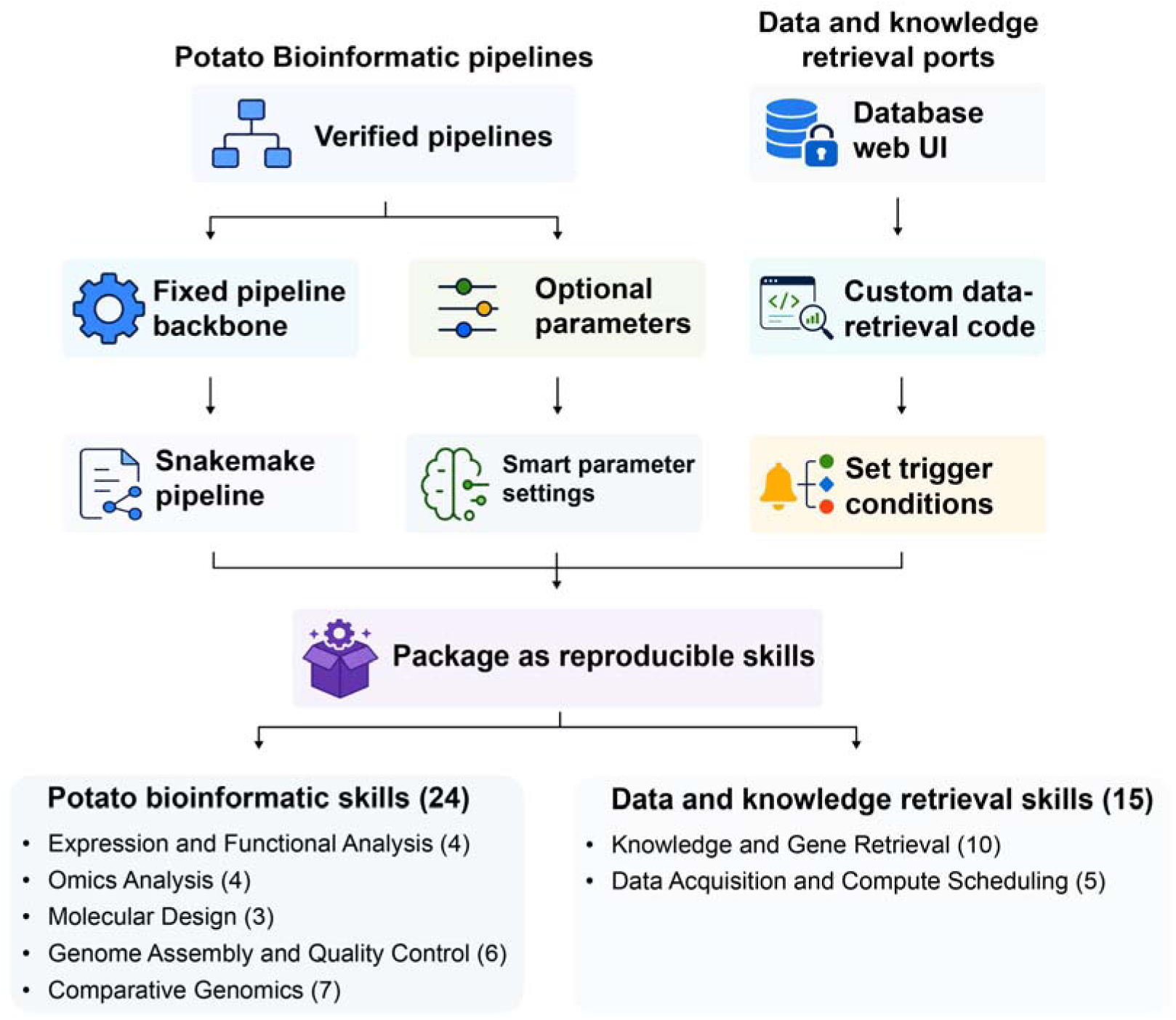
Development and organization of potato-specific Agent Skills. Validated bioinformatics procedures were implemented as fixed Snakemake workflow backbones with configurable parameters, whereas database access was implemented using custom retrieval code and trigger conditions. These components were packaged as 24 bioinformatics analysis skills and 15 data and knowledge retrieval skills.

To improve analytical consistency and reproducibility, we implemented the procedures underlying the bioinformatics analysis skills as pre-built and tested Snakemake workflows. The analysis steps, order of software execution, and key parameters are specified in the Snakefile, while software paths, input data, and computational resources such as CPU and memory remain configurable by the agent for the local environment. Conda environments and version locking are used to record and control software dependencies. During execution, the agent selects the skill, configures the inputs, and launches the workflow, while Snakemake manages task dependencies and execution. This mechanism reduces missing steps and unintended parameter variation that can arise when an agent writes an analysis pipeline from a short prompt, allowing the same inputs to be processed through a consistent workflow.

Among the 15 data and knowledge query skills, we developed structured tools for querying published literature, gene-function databases, and knowledge-graph resources, including NCBI PubMed, TAIR, RiceData, PlantConnectome, and PlantScience.ai (Lim et al., 2025; Reiser *et al*., 2024; Yu et al., 2026). These tools enable an agent to retrieve cross-species evidence related to a target gene or agronomic trait. We also reconstructed the data-access interface of the Potato Knowledge Hub so that an agent can query potato functional genes, associated publications, gene identifiers, and sequence information through a unified interface. Together with the potato multi-omics database APIs developed in this study, these skills provide access to the pangenome, tissue expression, co-expression, and spatial transcriptomic data. An agent can therefore integrate direct evidence from potato studies, homologous gene functions in model and crop species, and multi-omics evidence to support candidate gene screening and functional inference.

In summary, we developed and released 24 bioinformatics analysis skills and 15 data and knowledge query skills, establishing an execution layer that connects natural-language questions with data queries, knowledge retrieval, and reproducible analysis workflows. These openly available skills support researchers in conducting multi-omics data exploration, functional gene discovery, and molecular design more consistently, while enabling the research community to reuse, evaluate, and extend these analytical capabilities.

### Genome-wide gene function prediction and evidence grading

In our previous work, the Potato Knowledge Hub systematically curated reported potato functional genes and their associated publications. However, compared with model plants and major crops such as Arabidopsis, rice, and maize, relatively few potato genes have direct functional evidence, limiting large-scale candidate gene screening and favorable allele discovery. In contrast, the extensive functional studies available in other plant species provide a valuable basis for inferring potato gene functions through homology. However, such information is distributed across databases and publications, and its use requires sequence comparisons, database queries, literature searches, and evidence integration, making manual analysis laborious and time-consuming.

To systematically integrate this cross-species evidence, we combined the Potato Knowledge Hub, external database query skills, and the potato gene expression atlas developed in this study to establish a genome-wide gene function prediction workflow (Figure 3). For each potato gene, the workflow first queries its name, functional description, and associated publications in the Potato Knowledge Hub. A large language model then summarizes the relevant findings to generate a direct potato evidence summary. For Arabidopsis homologs, the workflow retrieves standardized gene names and functional annotations from TAIR, queries knowledge-graph information from PlantConnectome and relevant studies from PubMed, and uses a large language model to integrate the results into an Arabidopsis evidence summary. For rice and maize homologs, gene names and basic annotations are obtained from RiceData and MaizeGDB, respectively, and combined with PubMed literature to generate rice and maize evidence summaries. In addition to direct potato and homolog-based evidence, potato tissue expression patterns are included as supporting information to assess whether a predicted function is consistent with the tissues in which the gene is expressed.

**Figure 3.**
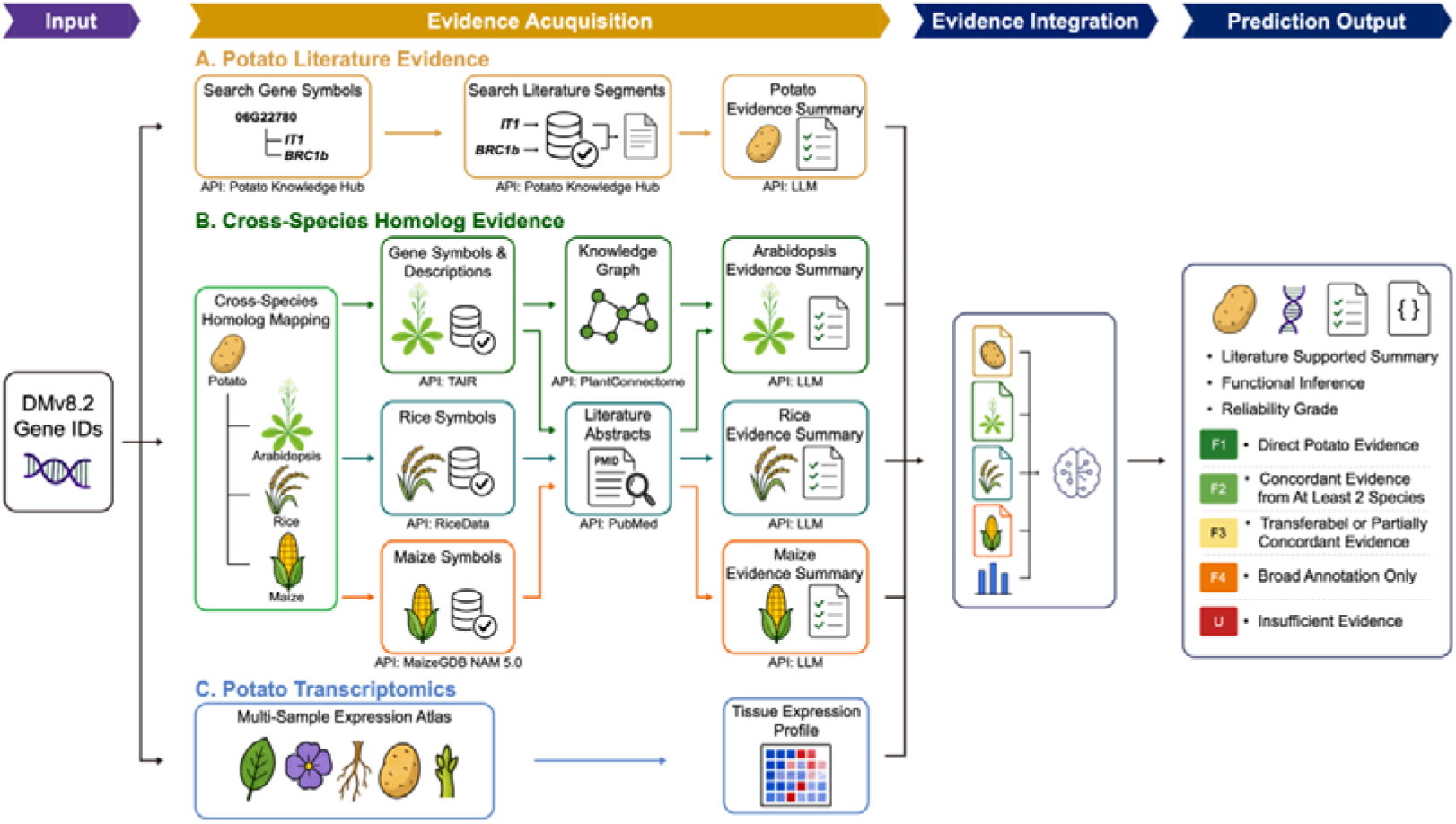
Workflow for genome-wide potato gene function prediction and evidence grading. DM v8.2 gene identifiers were used to collect (A) potato literature evidence from the Potato Knowledge Hub, (B) homolog-based evidence from Arabidopsis, rice, and maize through TAIR, PlantConnectome, RiceData, MaizeGDB, and PubMed, and (C) tissue expression evidence from the potato bulk RNA-seq atlas. Species-specific evidence was summarized by a large language model and integrated to generate an evidence summary, a putative functional assignment, and a reliability grade.

After evidence collection, direct potato evidence, homolog functions in Arabidopsis, rice, and maize, and potato tissue expression patterns are jointly provided to a large language model to generate an integrated functional overview and a putative functional assignment. To distinguish reported findings from computational inference, we established a predefined five-level evidence system comprising F1-F4 and U. F1 indicates direct functional evidence in potato supported by genetic and molecular experiments. F2 indicates consistent functional evidence for homologs in at least two species. F3 indicates that homologs have reported functions but that the evidence is less consistent among species. F4 indicates that only protein-domain or basic annotation information is currently available. U indicates that the collected evidence is insufficient to support a reliable functional prediction. This system allows users to distinguish the strength of evidence underlying each prediction and avoids treating cross-species inference as equivalent to experimentally validated gene function.

Using this workflow, we generated functional predictions and evidence levels for 37,658 genes in the DM v8.2 reference genome. Of these, 837 genes were classified as F1, accounting for 2.2% of all genes; 999 were classified as F2 (2.7%); 16,893 as F3 (44.9%); 11,752 as F4 (31.2%); and the remaining 7,177 as U (19.1%). The small proportion of F1 genes indicates that only a limited number of potato genes currently have direct functional evidence. In addition, more than half of all genes have only basic annotations or insufficient evidence, highlighting the gap between current potato functional genomics knowledge and the gene-level information required for potato breeding.

In summary, this workflow converts dispersed evidence from potato studies, cross-species homolog functions, and tissue expression into genome-wide, queryable, and graded functional evidence. It provides a systematic reference for AI-assisted candidate gene screening, functional gene discovery, and analysis of breeding traits. For streamlined access, the corresponding query script is wrapped in the potato-knowledge-search skill.

### Development of a multi-user Potato Agent

The potato multi-omics database, open APIs, Agent Skills, and genome-wide functional prediction results described above are available through open interfaces or as open-source resources and can be called by compatible, locally deployed agents. Personal agent deployment, however, still requires model access, runtime configuration, skill installation, dependency management, and suitable computing and storage resources. Standard personal computers are often insufficient for large-scale multi-omics data and computationally intensive bioinformatics analyses. To reduce this barrier, we developed Potato Agent as a multi-user platform for the potato research community, integrating the data, knowledge, skills, and analysis workflows described above into a directly accessible online service.

Potato Agent was developed from the open-source Hermes Agent framework (https://github.com/NousResearch/hermes-agent). We streamlined and reconstructed the original framework by removing instant messaging, personal dashboards, and other modules unrelated to scientific services, while retaining core capabilities such as the agent loop, tool and skill use, contextual memory, and sub-agent collaboration. We then added file-management and permission-based isolation modules for multi-user operation. Each Potato Agent account is assigned a dedicated workspace (Figure 4A), where users can manage their files and analysis jobs run under the corresponding user identity without access to other users’ data.

**Figure 4.**
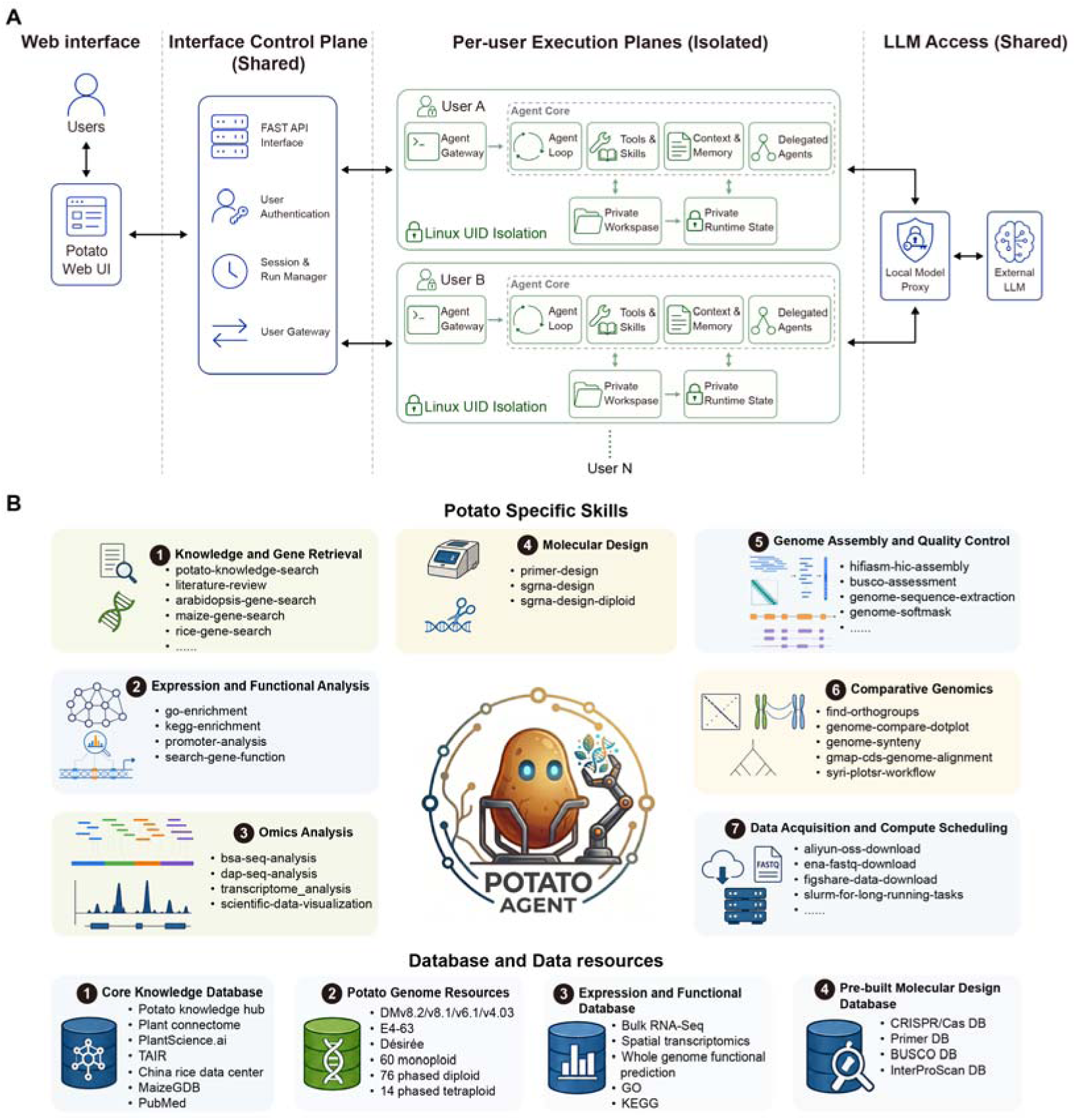
Architecture and integrated resources of Potato Agent. (A) Multi-user architecture of Potato Agent. A shared interface control plane handles authentication, session and run management, and request routing. Each account is mapped to a distinct Linux user identifier and an isolated execution environment containing a private workspace and running state. Each agent core provides the agent loop, tools and skills, context and memory, and delegated agents, while model requests are routed through a shared local proxy. (B) The 39 Agent Skills are organized into seven categories and connected to four classes of resources: core knowledge databases, potato genome resources, expression and functional databases, and molecular design and annotation databases.

To simplify interaction with the agent and facilitate inspection of analytical procedures and outputs, we developed the Potato Agent Web UI using FastAPI. The left panel manages previous chat and task histories, the central panel contains the conversation between the user and the agent, and the right panel displays the current user’s workspace (Supplementary Figure 3). Logs, tables, figures, sequences, and other result files generated during agent execution are saved in this workspace and can be previewed or downloaded through the web interface. This design allows users to inspect intermediate files and final outputs while interacting through natural language, improving the transparency and usability of agent-assisted analyses.

We integrated the 39 open-source Agent Skills described above into Potato Agent (Figure 4B). The platform also incorporates four categories of data resources. Core knowledge resources support queries for gene information, functional descriptions, and associated publications in potato, Arabidopsis, rice, and maize. Genome resources support queries of genomic features and extraction of accession-specific sequences. Expression and functional resources provide access to tissue expression patterns, co-expression relationships, genome-wide functional predictions, and GO and KEGG enrichment results. Molecular design and annotation services support CRISPR/Cas sgRNA design, PCR primer design, InterProScan domain annotation, and BUSCO completeness assessment. Through these resources and skills, Potato Agent translates natural-language questions into database queries, cross-species evidence integration, or reproducible bioinformatics tasks and returns the resulting files to the user’s workspace.

The Potato Agent source code and associated skills are available on GitHub and are being updated (https://github.com/biojiayuxin/potato-agent). Institutional users can deploy the platform on local servers or high-performance computing clusters, while individual users can access the online platform free of charge at https://potato-agent.ynnu.edu.cn.

In summary, Potato Agent integrates agent-ready multi-omics data, functional predictions, and open-source skills into a unified platform that supports online task execution, and result preview. The platform reduces the technical burden of deploying and using AI agents in potato research and provides researchers with a directly accessible environment for knowledge retrieval, data analysis, functional gene discovery, and results visualization.

### Case studies of Potato Agent

To assess Potato Agent in practical research tasks, we designed three representative case studies covering reproducible bioinformatics analysis, functional gene screening and validation, and haplotype-aware promoter analysis and sgRNA design. These cases examined whether Potato Agent could combine the databases, functional predictions, and open-source skills described above to translate scientific questions expressed in natural language into data queries, analysis workflows, and molecular design tasks.

#### 1 Reproducible bioinformatics analysis

We first used a published BSA-seq dataset for potato tuber flesh color to test Potato Agent on a complex bioinformatics workflow. BSA-seq analysis requires recognition of chromosome naming formats, coordinated filtering by sequencing depth and SNP depth, and ordered execution of shell, Python, and R scripts, making it suitable for evaluating an agent’s ability to configure and reuse a pre-built workflow. Following the user’s instruction, Potato Agent called the bsa-seq-analysis skill and completed data processing, variant filtering, statistical analysis, and result visualization.

The analysis detected a significant interval near the beginning of chromosome 12 that overlapped the previously reported *Yellow Leaf* locus (Figure 5A). The ΔSNP-index profile generated by Potato Agent was consistent with the BSA-seq profile reported in the previous study, including the major association signal on chromosome 12 (Li et al., 2024). These results show that Potato Agent can consistently execute a pre-built workflow and reproduce a previously reported mapping result.

**Figure 5.**
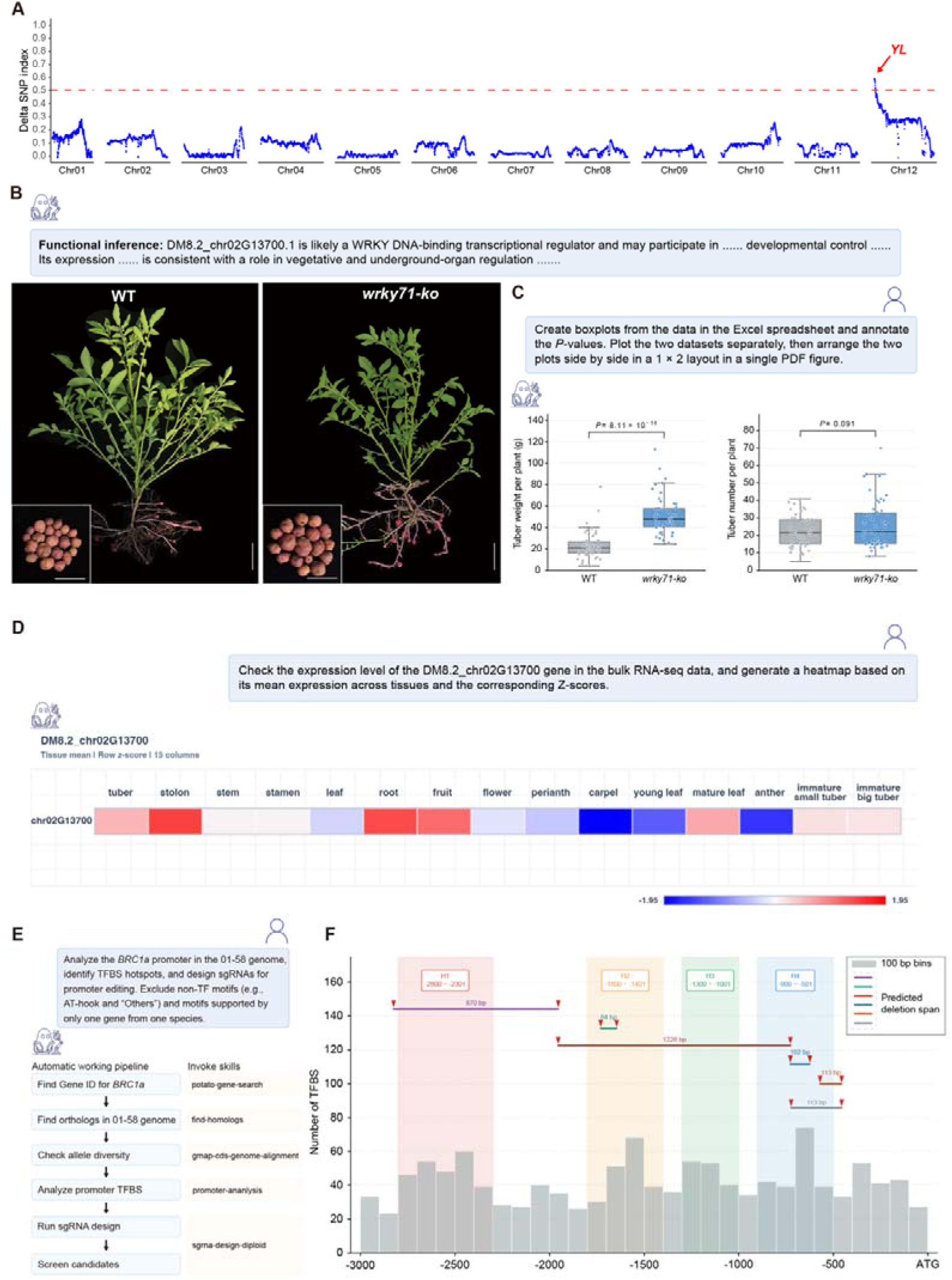
Case studies of Potato Agent. (A) Potato Agent executed a pre-built BSA-seq workflow and reproduced the previously reported ΔSNP-index profile, identifying the major association signal that overlapped the reported *YL* locus; the red dashed line marks a ΔSNP-index threshold of 0.5. (B) Functional inference for *DM8.2_chr02G13700* and representative phenotypes of wild-type (WT) and *wrky71* knockout plants at 60 days after planting. Insets show harvested tubers per plant. Scale bars: 4 cm. (C) Boxplots and individual observations for tuber weight and tuber number per plant. (D) Heatmap directly generated by Potato Agent from the bulk RNA-seq atlas in response to the user’s request, showing the Z-score-normalized mean expression of *WRKY71* across 15 tissue categories. (E) Potato Agent workflow and invoked skills for identifying *BRC1a* alleles, analyzing promoter transcription factor-binding sites (TFBSs), and designing and screening single-guide RNAs (sgRNAs). (F) Distribution of predicted TFBSs in 100-bp bins across the 3-kb *BRC1a* promoter. Shaded regions H1-H4 denote TFBS-dense regions; colored markers and connecting lines indicate six proposed editing combinations constructed from eight selected sgRNAs and their predicted deletion spans. ATG marks the translation start site.

#### 2 Agent-assisted identification and visualization of a tuber development regulator

We next examined whether the genome-wide functional predictions could be combined with Potato Agent for candidate gene screening and functional evidence analysis. In our previous study, multi-omics comparisons identified 28 transcription factors that may regulate tuber development. Among them, *Identity of Tuber1* (*DM8.2_chr06G22780*) was shown to be a tuber identity gene that controls tuber initiation (Tang *et al*., 2022). To identify additional regulators from the remaining candidate transcription factors, we combined this candidate set with the genome-wide functional prediction results and selected *WRKY71* (*DM8.2_chr02G13700*) for validation. Cross-species homolog evidence suggested that this gene may be involved in underground organ development (Figure 5B, https://potato-agent.ynnu.edu.cn/genes/DM8.2_chr02G13700).

We knocked out *WRKY71* in potato and used Potato Agent to perform statistical tests and visualize total tuber weight per plant and tuber number in wild-type and *wrky71* mutant plants (Figure 5C, Supplementary Figure 4A). Loss of *WRKY71* significantly increased tuber weight per plant but did not significantly affect tuber number, suggesting that *WRKY71* negatively regulate tuber yield (Figure 5C). We then asked Potato Agent to retrieve the mean expression of *WRKY71* across tissues from the bulk RNA-seq atlas and visualize the values after Z-score normalization (Figure 5D). *WRKY71* showed relatively high expression in tubers and stolons, consistent with a possible role in tuber development.

To examine genomic conservation around *WRKY71*, we asked Potato Agent to retrieve genes within 100 kb upstream and downstream of *WRKY71*, compare local synteny between the DM and E4-63 genomes, and generate a visualization. Potato Agent extracted the neighboring genes, organized homologous relationships, and constructed the local synteny plot from the underlying genome resources (Supplementary Figure 4B). The database did not contain a precomputed *WRKY71* flanking-gene set or a gene-specific DM-E4-63 synteny result; the agent generated both in response to the user’s request. This case therefore connected genome-wide functional prediction, experimental phenotype statistics, tissue expression queries, and a user-defined comparative genomic analysis in a continuous workflow, illustrating the autonomy of Potato Agent in retrieving and combining data for candidate gene analysis.

#### 3 Haplotype-aware promoter analysis and sgRNA design

Promoter editing can modulate gene expression without altering protein-coding sequences and offers a potential strategy for improving crop architecture. For example, editing the promoter of rice *IPA1* can adjust the balance between grain number per panicle and tiller number, thereby increasing grain yield per plant (Song et al., 2022). Cultivated potato tends to develop excessive branches late in the growing season, which can increase non-productive biomass and complicate field management. *BRC1* has a conserved role in axillary branch development, and its potato homolog *BRC1a* regulates branching of both shoots and stolons (Nicolas et al., 2015). Therefore, this gene can be a potential target for improving potato architecture through promoter editing.

Promoter editing in a highly heterozygous potato accession requires consideration of promoter sequence differences between haplotypes, the distribution of transcription factor-binding sites (TFBSs), sgRNA targeting efficiency, and off-target risk. We therefore developed the promoter-analysis and sgrna-design-diploid skills based on the PlantPAN 4.0 web API (https://plantpan.itps.ncku.edu.tw/plantpan4/) and CRISPOR software, respectively, and used *BRC1a* to evaluate Potato Agent’s ability to perform haplotype-aware promoter analysis and sgRNA design (Figure 5E, F).

Potato Agent first extracted and compared the 3-kb sequence upstream of *BRC1a* from each of the two 01-58 haplotype assemblies. The two promoter sequences were identical across this region, indicating that the same sgRNAs could be used to target both *BRC1a* alleles. Potato Agent then performed cis-regulatory element analysis of the *BRC1a* promoter and identified 1,255 predicted TFBSs. Based on their distribution along the promoter, four TFBS-dense regions were identified. These regions contained 247, 188, 147, and 194 predicted sites, respectively, accounting for 776 sites in total. Potato Agent subsequently designed sgRNAs for these regions and filtered them by predicted off-target risk, target-sequence features, and editing efficiency, yielding 28 candidate sgRNAs. Potato Agent then selected eight targets with relatively high predicted editing efficiencies and arranged them into six promoter-editing combinations. These combinations were designed to cover different TFBS-dense regions and to support experimental testing of promoter variants with potentially different regulatory effects (Figure 5E, F).

Together, the three cases illustrate the feasibility and practical utility of Potato Agent at three levels: recovery of a known locus, functional gene discovery, and molecular design. Potato Agent can combine standardized multi-omics data, genome-wide functional predictions, and reproducible analysis skills to support a continuous process from scientific question formulation and evidence retrieval to data analysis, candidate gene screening, and experimental design. These examples demonstrate how a unified agent environment can support functional gene discovery and molecular breeding research in potato.

## Discussions

In response to the need for functional gene discovery and favorable allele stacking in potato breeding, we developed a data, knowledge, and analysis framework for AI agents. We integrated genome resources from 150 potato accessions, 259 bulk RNA-seq samples, and 14 spatial transcriptomic datasets to construct a pangenome, a tissue expression atlas, co-expression networks, and spatial expression maps. We also generated functional predictions and evidence levels for 37,658 genes in the DM v8.2 reference genome. On this basis, we developed and released 24 bioinformatics analysis skills and 15 data and knowledge query skills and incorporated these resources into Potato Agent, which supports multi-user data separation and online result preview. Case studies involving BSA-seq analysis, identification of a tuber development regulator, and promoter analysis and sgRNA design, further illustrated the utility of this system for data analysis, knowledge integration, and molecular design.

Existing potato databases provide an important basis for genomic research and functional gene retrieval, but different platforms generally emphasize genome data, functional genes, or literature knowledge separately. Unified integration of pangenomic, transcriptomic, spatial transcriptomic, and functional evidence remains limited. Conventional databases also commonly rely on query pages, datasets, and analysis modules configured in advance by developers. Analyses are therefore often constrained by the available functions, making it difficult to flexibly select data, adjust parameters, or combine analytical steps for a specific scientific question. The platform developed here integrates potato multi-omics data and provides complementary access for researchers and AI agents. Researchers can query and browse data through the web interface, while agents can retrieve structured information through open APIs and call Agent Skills for knowledge retrieval and data analysis. This design extends the database from a resource primarily used for information access to an analytical infrastructure whose components can be called and combined by agents.

Potato Agent extends the Potato Research Assistant previously developed within the Potato Knowledge Hub. Whereas the earlier assistant focused on literature and functional gene retrieval, Potato Agent integrates multi-omics data, cross-species evidence, bioinformatics workflows, and molecular design. Its multi-user workspaces and online result preview further enable it to process user data and execute computational tasks.

The pre-built skills improve the consistency and reproducibility of common analyses, but they do not define the functional boundary of Potato Agent. For tasks with established procedures, such as BSA-seq, Potato Agent can directly call validated Snakemake workflows. For questions not yet covered by a skill, the agent can combine database interfaces, command-line tools, and existing skills, write task-specific analysis code, and use sub-agents to decompose complex tasks. New workflows that have been repeatedly tested can subsequently be packaged as open-source skills. Potato Agent can therefore support consistent analyses through standardized skills while remaining extensible to new research questions.

The current platform mainly integrates genomic, transcriptomic, and functional knowledge resources and does not yet systematically include genotype and phenotype data from natural populations. Although germplasm repositories such as International Potato Center have curated some phenotype information, agronomic traits in potato are strongly influenced by geography, climate, and cultivation conditions, and many phenotypic datasets with value for local adaptation remain insufficiently standardized and shared. In the future, we aim to work with the potato research and breeding community to standardize and integrate genotypic and phenotypic data for potato germplasm and develop a shared data platform for gene discovery and hybrid breeding.

## Methods

### Pangenome analysis

Protein sequences from 135 diploid potato genomes were used for gene-family (orthogroup) inference. To avoid the redundant allelic genes in phased diploid genomes, only one haplotype was used for analysis. OrthoFinder v3.1.5 was ran with parameters -b, -og, --old-version, to category gene-families (Emms et al., 2026). Gene families were classified into mutually exclusive categories based on their occurrence across the 135 genomes: core families were present in all genomes; soft-core families were present in more than 90% but fewer than all genomes (122–134 genomes); dispensable families were present in 2–121 genomes; and private families were present in only one genome.

Pan- and core-genome accumulation curves were generated from the binary orthogroup-presence matrix. For each number of sampled genomes, the expected numbers of pan and strict-core gene families were calculated from the orthogroup occupancy distribution using hypergeometric probabilities. In addition, 1,000 nested random genome-order permutations were performed using a fixed random seed (20260725), and the 2.5%, median, and 97.5% quantiles were used to display order-dependent variation. Endpoint-anchored biexponential curves were fitted descriptively to the expected pan- and core-genome trajectories using SciPy, with parameter variation assessed from 200 randomly selected permutation trajectories.

### RNA-Seq analysis

Raw reads were adapter-trimmed and quality-filtered with fastp v1.3.3 using automatic adapter detection and default filtering settings (Chen, 2023). Clean reads were aligned to the DM v8.2 reference genome using HISAT2 v2.2.2 with the --dta option (Kim et al., 2019). Alignments were converted to BAM format, coordinate-sorted, and indexed using SAMtools v1.23.1 (Li et al., 2009). Annotation-guided expression quantification was performed against the DM v8.2 GFF3 annotation using StringTie v3.0.3 with the parameters -e -B -A -p 8 -G (Shinder et al., 2026).

### Weighted Gene Co-expression Network Analysis

Samples were divided into five tissue/developmental groups: leaf (n = 57), stem (n = 53), root (n = 51), reproductive tissues (n = 49), and tuberization-related tissues (n = 49). Within each group, genes with TPM ≥ 1 in at least 20% of samples were retained and transformed as log (TPM + 1). The 12,000 genes with the highest expression variance were used for network construction.

Weighted gene co-expression networks were constructed in R 4.5.3 using the WGCNA package v1.74 (Langfelder and Horvath, 2008). Biweight midcorrelation (bicor) was used to generate signed adjacency matrices and signed topological overlap matrices. Soft-thresholding powers were selected as the lowest values achieving a signed scale-free topology fit of R² ≥ 0.80 and were 10, 6, 14, 11, and 10 for the leaf, stem, root, reproductive, and tuberization networks, respectively. Modules were identified using blockwiseModules with a minimum module size of 30, reassignThreshold = 0, pamRespectsDendro = FALSE, and mergeCutHeight = 0.25. Module eigengenes and gene module membership (kME) were subsequently calculated using biweight midcorrelation.

### Genome assembly

PacBio HiFi reads and Hi-C data were used for genome assembly. The 19-mer frequency distribution was calculated using KAT v2.4.2, and genome size and heterozygosity were estimated using GenomeScope 2.0 (Mapleson et al., 2017; Ranallo-Benavidez et al., 2020). Haplotype phasing and initial assembly were performed with hifiasm v0.25.0-r726 using the Hi-C data with parameters --hg-size 800m --hom-cov 44 --primary (Cheng et al., 2022).

For each haplotype, a genome database was first constructed using BuildDatabase, a utility included in the RepeatModeler v2.0.4 package, with the parameters -name repeatdatabase -engine ncbi (Flynn et al., 2020). A haplotype-specific *de novo* repeat library was then generated using RepeatModeler. Each genome was soft-masked with RepeatMasker v4.1.5 using its corresponding de novo repeat library (https://www.repeatmasker.org/). Paired-end RNA-seq libraries derived from tuber, leaf, stem, stolon, and axillary bud tissues were aligned separately to each soft-masked assembly using HISAT2 v2.2.1 with the --dta option (Kim *et al*., 2019). Reference proteomes from potato DM v6.1 and DM v8.2 and tomato Heinz 1706 SL4.0 were used as homologous protein evidence. Gene models were predicted independently for each haplotype using BRAKER3 v3.0.8 in ETP mode, with the soft-masked genome assembly, RNA-seq alignment files, and combined homologous protein sequences supplied using --genome, --bam, and --prot_seq (Gabriel et al., 2024).

Genome assembly and annotation completeness was evaluated using BUSCO v6.1.0 against the solanales_odb12 lineage dataset, using genome mode (-m genome) for the assemblies and protein mode (-m proteins) for the representative protein sets (Tegenfeldt et al., 2025).

### Genome comparison

Chromosome-level whole-genome comparison was performed using reference and query genome assemblies in FASTA format. The query assembly was aligned to the reference assembly using nucmer from MUMmer4 v4.0.1 with the parameters -l 100-c 500 (Marcais et al., 2018). One-to-one alignments were retained using delta-filter with the parameters −1 -i 90 -l 1000. The filtered alignments were converted into tab-delimited coordinates using show-coords with the options-THrd. Syntenic regions and structural rearrangements were identified using SyRI v1.7.1 with the filtered delta and coordinate files as input and the --nosnp option to exclude SNP and small-indel calling (Goel et al., 2019). The resulting syntenic relationships and structural rearrangements were visualized using plotsr v1.1.1 (Goel and Schneeberger, 2022).

Macroscopic and microscopic synteny analyses were performed using the Python implementation of MCScan in JCVI v1.6.5 (Tang et al., 2024). Genome annotation files in GFF3 format were converted to BED format, and the corresponding protein sequences were used for homology searches with DIAMOND BLASTP v2.1.25 (Buchfink et al., 2021). Syntenic gene pairs and collinear blocks were identified using jcvi.compara.catalog ortholog with --cscore=0.9 and a minimum block span of 30 genes. The resulting anchor files were simplified into chromosome-level collinear blocks and visualized as macrosynteny plots using jcvi.graphics.karyotype. For microsynteny analysis, local collinear regions were extracted from the lifted anchor file using jcvi.compara.synteny mcscan with --iter=1, which retained the best matching region for each reference interval. Gene order and homologous relationships within selected genomic intervals were visualized using jcvi.graphics.synteny with arrow-shaped gene models and orthogroup-based coloring.

### Identify orthologs among potato genomes

Orthologs between closely related potato genomes were identified using MCScan in JCVI package (v1.6.5), with parameters --dbtype=prot, --cscore=0.7,--no_strip_names, and --no_dotplot. Ortholog assignments were preferentially selected from the resulting ‘.anchors’ file using the highest-scoring anchor pairs. For genes lacking anchor-supported matches, hits from ‘.last.filtered’ with ≥90% sequence identity were considered, and the candidate with the highest bit score was retained.

### Identify potato homologs in arabidopsis and crops

Cross-family potato homologs were identified using a three-level workflow combining JCVI/MCScan synteny and BLASTP sequence similarity. Collinear gene pairs were detected using jcvi.compara.catalog ortholog with --dbtype=prot, --align_soft=blast, --cscore=0.7, --min_size=4, and --no_strip_names, and the highest-scoring anchor pair for each potato gene was retained as the primary homolog (L1). For genes without synteny-supported matches, bidirectional BLASTP searches were performed using an E-value cutoff of 1e-5, -max_target_seqs 20, and -max_hsps 1; reciprocal best hits were classified as L2, while the remaining potato genes were assigned their best significant target-species BLASTP hit as L3.

### GO and KEGG enrichment analysis

Gene Ontology (GO) annotations were retrieved in OBO format from the OBO Foundry (http://purl.obolibrary.org/obo/go/go-basic.obo), while KEGG pathway annotations were extracted from the built-in TBtools.KeggBackEnd database in TBtools-II (Chen et al., 2023). The gene annotation file for the DMv8.2 genome was generated using the online platform eggNOG-mapper (http://eggnog-mapper.embl.de/) with default parameters. After establishing the correspondence between the annotation file and the background files, functional enrichment analysis was performed using the enricher function from the R package ClusterProfiler (v4.12.6), with the significance threshold set to pvalueCutoff = 0.05 and adjusted using the Benjamini–Hochberg correction (Wu et al., 2021). The enrichment results were visualized using the ggplot2 package (v3.5.2) (Wickham, 2016).

### sgRNA design

The genome assembly (FASTA) and corresponding gene annotation (GFF3) were used to construct a genome-specific database with CRISPOR and BWA v0.7.19-r1273 (Concordet and Haeussler, 2018; Vasimuddin et al., 2019). The genome was converted to 2bit format, indexed using BWA, and integrated with gene annotations using crispor-add-genome. sgRNAs were designed using CRISPOR for SpCas9, with a 20-nt guide sequence, an NGG PAM, and genome-wide off-target searches allowing up to four mismatches (-p NGG --guideLen 20 --mm 4). Candidate sgRNAs were ranked based on MIT and CFD specificity scores and predicted on-target efficiency.

### PCR primer design and in silico specificity assessment

Primer pairs were designed using primer3-py v2.3.0 (Untergasser et al., 2012). Parameters were set as follows: primer length, 18–25 nt (optimum, 20 nt); Tm, 58–62°C (optimum, 60°C); GC content, 40–60%; maximum pairwise Tm difference, 2°C; 3′ GC clamp, 1 nt; and maximum homopolymer length, 4 nt. Twenty candidate primer pairs were generated for each target, and the desired amplicon-size range was specified according to the experimental requirements. Hairpin formation, self-dimerization, heterodimerization, and 3′-end stability were evaluated using the thermodynamic functions implemented in primer3-py. Primer specificity was subsequently assessed by pair-wise in silico PCR against the corresponding reference genome using MFEprimer v4.2.4 (Wang et al., 2019). The genome database was indexed using a k-mer size of 9, and the amplicon-size limits used for specificity screening were adjusted to match the predefined experimental requirements. Primer pairs producing the expected target amplicon without additional credible products within the specified size range were retained for experimental validation.

### BSA-Seq analysis

Raw paired-end reads from the two contrasting bulks were quality-filtered using fastp v1.3.3 with automatic paired-end adapter detection (Chen, 2023). Clean reads were aligned to the reference genome using BWA-MEM v0.7.19-r1273, with sample-specific Illumina read-group tags, and alignments were sorted and indexed using SAMtools v1.23.1 (Li *et al*., 2009; Vasimuddin *et al*., 2019). PCR duplicates were marked using GATK MarkDuplicates v4.6.2.0 (McKenna et al., 2010). Variants were called independently for each bulk and chromosome using BCFtools (v1.23.1) mpileup/call (mpileup -Ou -r chrom -f reference; call -m -V indels) (Danecek et al., 2021). Variants were filtered by QUAL > 50 and by sample-specific sequencing depth, defined as one-third to threefold of the estimated bulk depth derived from total FASTQ bases. For each retained SNP, the SNP-index was calculated as the alternate-allele depth divided by total allele depth. ΔSNP-index was calculated as SNP-index_bulk A − SNP-index_bulk B. Genome-wide signals were summarized in sliding windows of 1 Mb with a 100-kb step. The absolute mean ΔSNP-index was plotted across chromosomes, and windows with ΔSNP-index values above 0.4 were highlighted as candidate associated regions.

### Development of Potato Agent

Potato Agent was developed as a browser-based application. The frontend was implemented using HTML5, CSS, and JavaScript. The backend was developed in Python using the FastAPI framework and deployed with Uvicorn. The frontend communicates with the backend through RESTful APIs using JSON over HTTPS, while real-time agent responses are transmitted through authenticated WebSocket connections. The backend communicates with each user-specific Hermes Lite runtime using JSON-RPC 2.0. Application data, including user accounts, conversations, runtime states, and archives, are primarily stored in SQLite relational databases and queried using SQL.

## Acknowledgements

This work was supported by National Natural Science Foundation of China (32488302, 32302582, 32402608, U2202206, 32361143517), Guangdong Major Project of Basic and Applied Basic Research (2021B0301030004), Agricultural Science and Technology Innovation Program (CAAS-ZDRW202404), Yunnan Fundamental Research Projects (202501BC070003, 202501AT070005), Yunnan Province Xingdian Talent Support Plan (XDYC-YLXZ-2022-0019).

## Declaration of interests

The authors declare no competing interests.

## Author contributions

S.H., Y.S., Yuxin J. and Y.Z. conceived and designed the study. Yuxin J. and Yudong J. designed the architecture and wrote the codes of Potato Agent. Yuxin J. and Y.D. designed the user interface and performed comprehensive functional tests of Potato Agent. Yuxin J., Jinye L., Jilin L. performed the bioinformatic analysis. F.L., L.W., X.S. and J.H. performed transgenic verification of *WRKY71*. D.L. performed stability tests of Potato Agent. Yuxin J. and Y.Z. wrote the manuscript. S.H., Y.S., Yuxin J. and Y.Z. contribute to the discussion and revised the manuscript. All authors read and approved the final manuscript.

## Data and code availability

The genome data was obtained from SolOmics database (http://solomics.agis.org.cn) (Bao et al., 2022; Cheng *et al*., 2025; Tang *et al*., 2022; Zhang *et al*., 2025), FigShare (https://doi.org/10.6084/m9.figshare.28531079), Bioinformatics lab website (http://www.bioinformaticslab.cn/pubs/dm8/), Spud DB (https://spuddb.uga.edu/), GitHub (https://github.com/HongboDoll/PotatoNLRome/releases/tag/v1.2.1), Zenodo (https://doi.org/10.5281/zenodo.15282553, https://doi.org/10.5281/zenodo.14053896) (Coronejo et al., 2026; Dong et al., 2024; Godec et al., 2025; Sun et al., 2025; Wang et al., 2026; Yang et al., 2023; Zhu et al., 2025).

The bulk RNA-Seq data was obtained from NCBI SRA under BioProject PRJNA754534, PRJNA573826, and PRJNA1173794 (Tang *et al*., 2022; Zhang *et al*., 2025; Zhou et al., 2020). The potato spatial transcriptomic data was obtained from NCBI SRA under BioProject PRJNA1462399.

The source code of Potato Agent was maintained at GitHub (https://github.com/biojiayuxin/potato-agent).

## Supplementary Figures

**Supplementary Figure 1.**
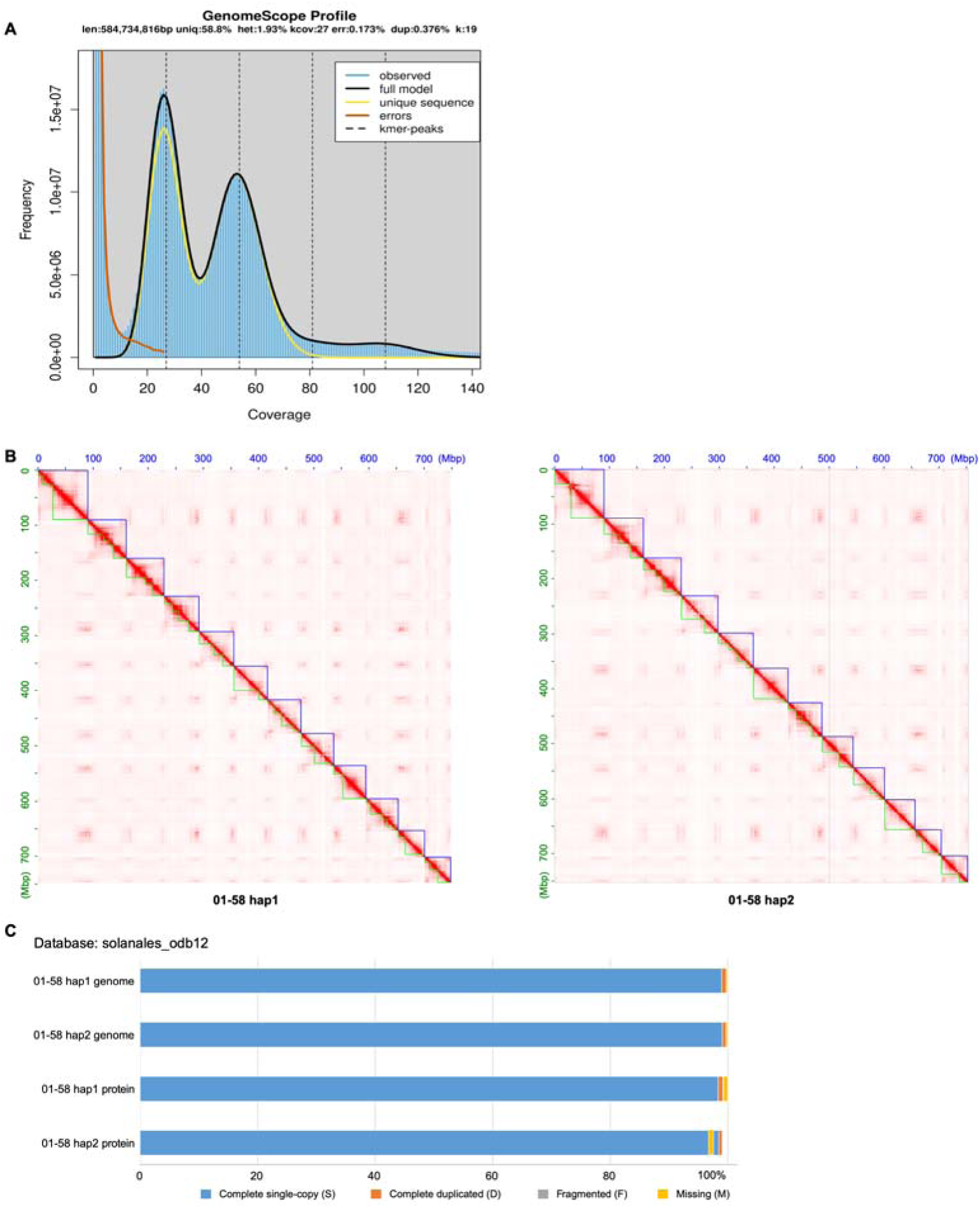
Genome assembly and annotation of diploid potato accession 01-58. (A) GenomeScope profile of 19-mer frequencies used to estimate genome size and heterozygosity. (B) Hi-C contact maps for haplotypes 1 and 2. (C) BUSCO assessment of the completeness of the two haplotype assemblies and their predicted gene models.

**Supplementary Figure 2.**
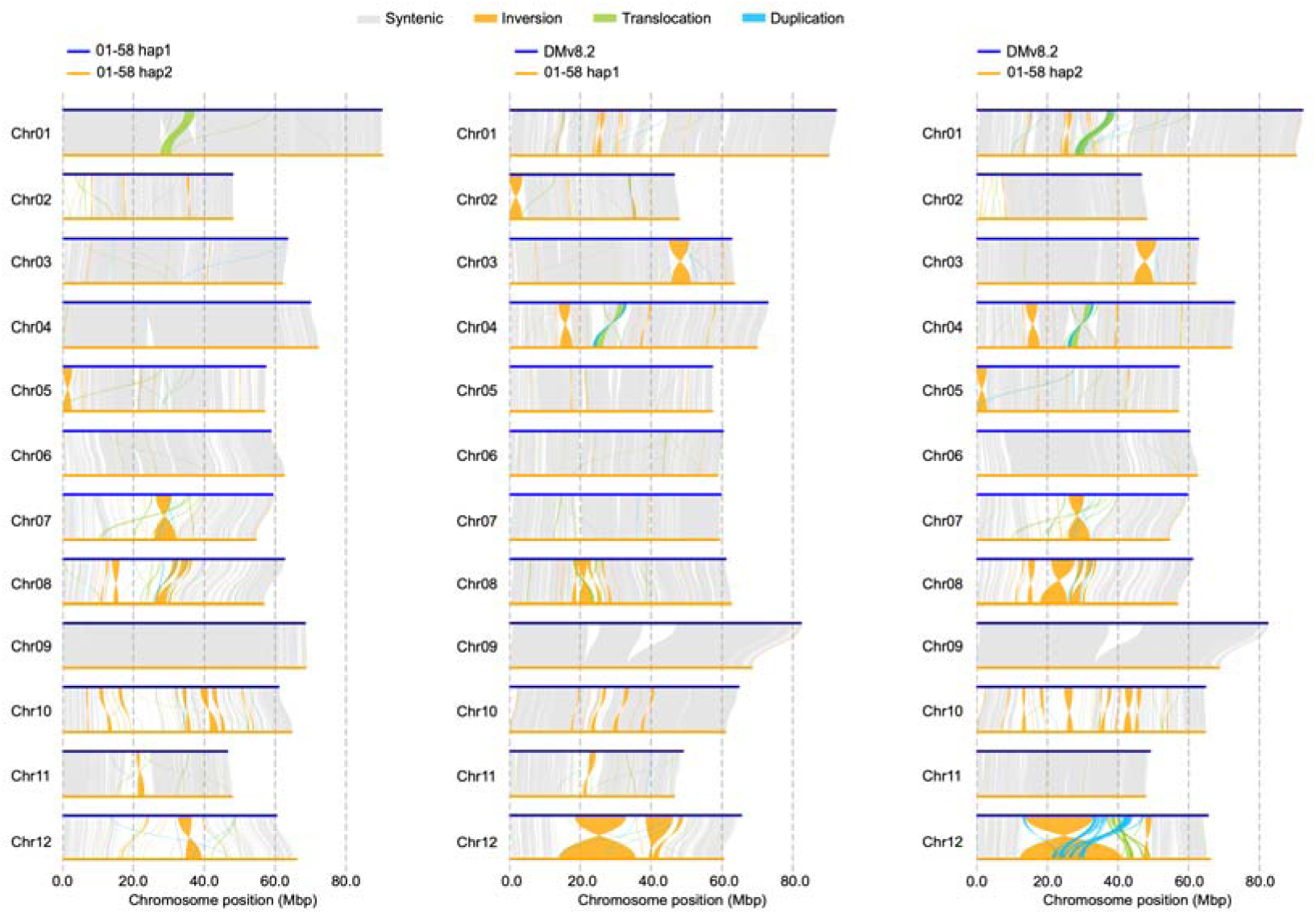
Whole-genome comparison of the 01-58 haplotypes and the DM v8.2 reference genome. Comparisons are shown between haplotype 1 and haplotype 2 (left), DM v8.2 and haplotype 1 (middle), and DM v8.2 and haplotype 2 (right). Gray, orange, green, and blue links denote syntenic regions, inversions, translocations, and duplications, respectively.

**Supplementary Figure 3.**
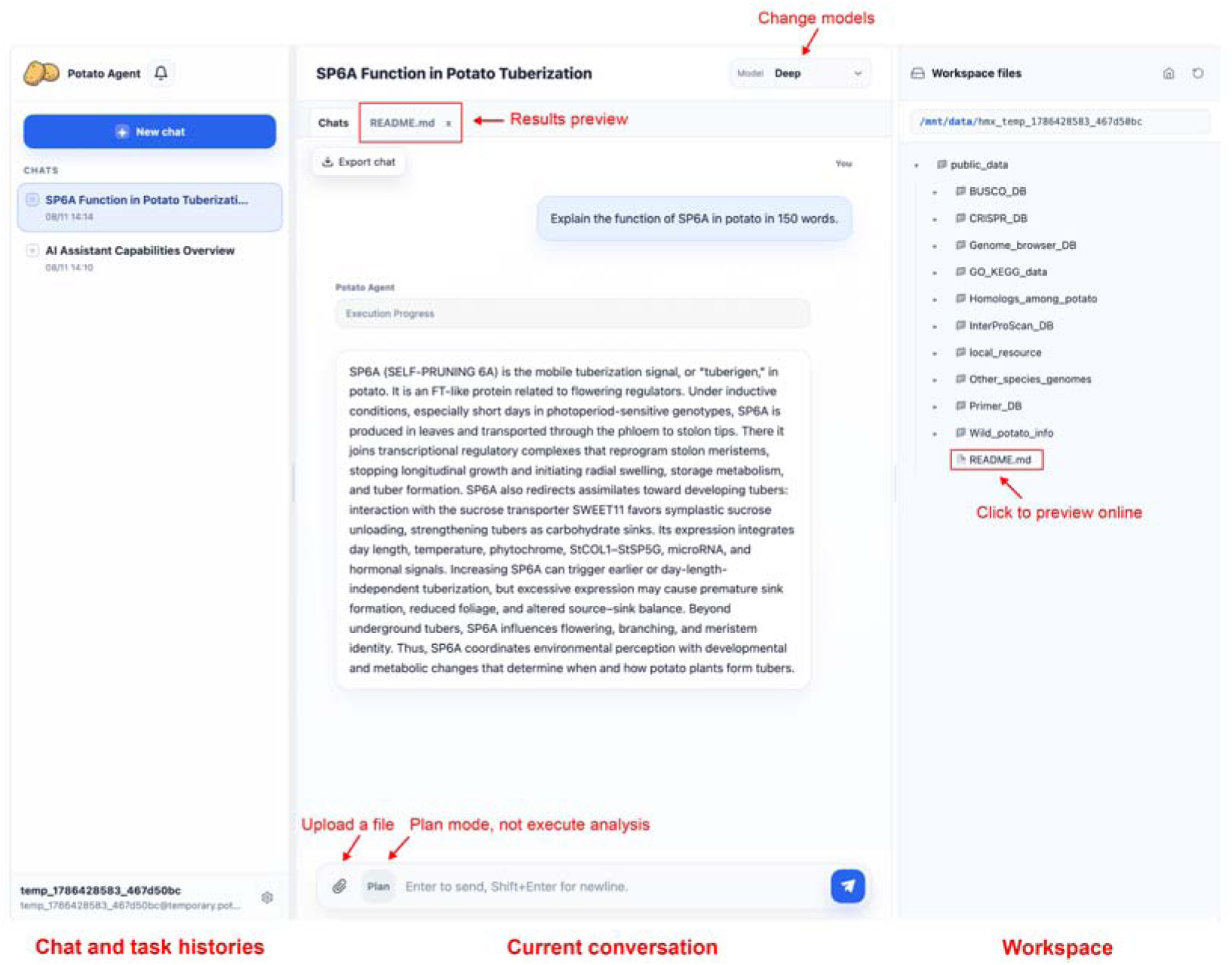
Web interface of Potato Agent. The left panel shows chat and task histories, the center panel displays the current conversation and result previews, and the right panel shows the user’s workspace and file tree.

**Supplementary Figure 4.**
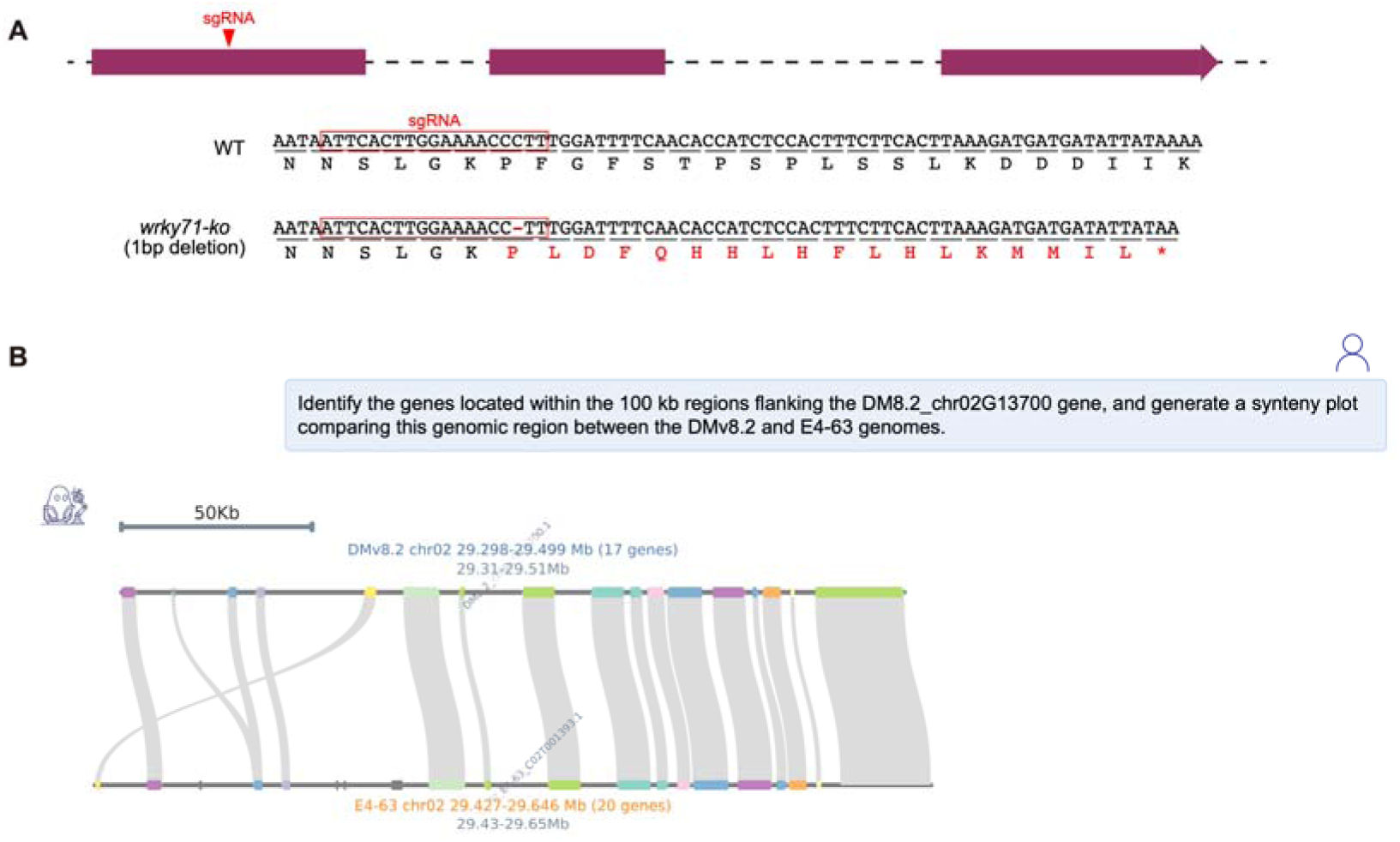
WRKY71 gene editing and local synteny analysis. (A) Schematic of *WRKY71* gene editing. The *wrky71-ko* allele contains a 1-bp deletion at the sgRNA target site, causing a frameshift and premature stop codon. (B) Microsynteny between the DMv8.2 and E4-63 genomic regions extending 100 kb upstream and downstream of the respective *WRKY71* loci, generated by Potato Agent in response to the user’s request.

## Notes

https://potato-agent.ynnu.edu.cn/

### Competing Interest Statement

The authors have declared no competing interest.

